# Postnatal Autonomic Development Predicts Adolescent Psychosocial Outcome

**DOI:** 10.64898/2026.08.03.742419

**Authors:** Maximilian Schmausser, Leonie Fleck, Anna Fuchs, Eva Möhler, Michael Kaess, Julian Koenig

## Abstract

**Background:** The maturation of the autonomic nervous system (ANS) has been suggested to play a crucial role in the development of emotion regulation and later psychosocial functioning. However, longitudinal evidence linking early autonomic development to long-term outcomes remains limited. This longitudinal study investigated the interplay between birth-related factors, early autonomic activity, and psychosocial outcomes across development.

**Methods:** The sample comprised 101 participants followed from two weeks to 14 years of age, with heart rate (HR) and vagally mediated heart rate variability (vmHRV) assessed at 2 weeks, 6 weeks, 3 months, 14 months, and 14 years. Linear models were used to examine associations between birth-related factors and early HR and vmHRV, as well as whether HR and vmHRV trajectories during the first 14 months predicted psychosocial outcomes at 5 and 14 years.

**Results:** Multiple birth-related factors significantly predicted HR and vmHRV at two weeks after birth. Moreover, flatter age-related increases in vmHRV and weaker decreases in HR during infancy predicted higher maternally reported psychosocial difficulties at 14 years in males only, with no such effects at 5 years or in females.

**Conclusions:** These findings underscore the importance of early autonomic maturation in shaping later psychosocial functioning, with effects on adolescent outcomes observed in males only. Early ANS trajectories may represent meaningful predictors of neurodevelopmental outcomes, highlighting their potential relevance for early identification of later psychosocial risk in a sex-specific manner.

## Introduction

Research over the past three decades has repeatedly demonstrated that the regulation of cardiac activity via the vagus nerve is a key component in the organism’s adaptation to environmental demands. Vagally mediated heart rate variability (vmHRV), referring to the temporal variations in beat-to-beat intervals between consecutive heartbeats, modulated by the vagus nerve, ^1,2^ has been associated with a wide range of physical and mental outcomes. Higher vmHRV has been linked to better cognitive functioning ^3^ and emotion regulation ^4^ whereas low levels of vmHRV represent a transdiagnostic marker for increased vulnerability to a variety of mental disorders, both in adulthood and adolescence ^5–7^.

Postnatal changes in vmHRV follow an characteristic developmental trajectory, with an early increase peaking during puberty, ^8–10^ followed by a gradual decline across adulthood, eventually reaching a lower plateau around the age of 60 years. ^11^ Despite this general pattern, the individual course of this trajectory underlies various physical, ^9^ psychological, ^12^ and environmental ^13^ influences. However, existing evidence predominantly relies on cross-sectional comparisons of different age groups, thereby largely failing to capture intra-individual variances across development.

Under a developmental neurovisceral framework, we recently suggested a close interconnection between autonomic nervous system (ANS) maturation, structural and functional brain development, and psychological functioning. ^14^ Accordingly, normative increases in vagal activity during early development mirror normative brain development, thus supporting cognitive, emotional and physiological functioning. Consequently, developmental influences impairing normative maturation of the ANS may directly impede both the development of the brain and the acquisition of cognitive and emotional skills. Supporting these assumptions, early life stress has been associated with decreased vmHRV, ^15,16^ perturbations in cognitive and emotional brain networks, ^17,18^ and elevated susceptibility to psychopathology. ^19^ McLaughlin et al. ^20^ further demonstrated that vmHRV moderates the association between psychosocial stress and psychopathology, such that psychosocial stress was positively related to internalizing symptoms in adolescents with low vmHRV but not in those with high vmHRV, underscoring the role of vagal activity in the development of psychiatric disorders. However, longitudinal studies, crucial to inform our understanding of developmental trajectories, are scarce. Most existing research focuses on adolescence and beyond, often examining treatment-related changes in vmHRV in association with symptom progression rather than developmental trajectories ^21–24^. Addressing this gap, we propose that early-life trajectories of vmHRV may offer valuable insights into psychopathological development.

Here we employed a within-subject longitudinal design spanning - to the best of our knowledge - the most extensive duration within this research area to date, assessing vmHRV and heart rate (HR) at six time points from two weeks to 14 years of age. We investigated the relationships between birth-related factors, maternal bonding, the early developmental trajectory of HR and vmHRV in infancy, and the emergence of psychopathological symptoms in childhood and adolescence. Specifically, we examined how prenatal factors, such as maternal stress and pregnancy complications, as well as impairments in maternal bonding impact vmHRV and HR in early infancy. We hypothesized that adverse birth-related conditions will be reflected in altered vmHRV and HR profiles, both through reduced baseline levels at the age of 2 weeks as well as through attenuated maturational change during infancy.

Additionally, based on our developmental neurovisceral framework^14^, we proposed that such deviations in autonomic developmental trajectories constitute early biomarkers of atypical psychosocial development, thereby conferring increased risk for the emergence of emotional and behavioral problems at ages 5 and 14 years.

## Methods & Materials

### Procedure

The study was approved by the Ethics Committee of the Faculty of Medicine at the University of Heidelberg (S-553/2016). The cohort was part of a longitudinal study, initiated in 2002/2003, two weeks after childbirth. The study involved six repeated assessments: 2 weeks after birth (T1), 6 weeks after birth (T2), at 4 months (T3), at 14 months (T4), at 5.5 years (T5), and at 14 years (T6) of offspring age, conducted between the 2002 and 2017. Mothers provided written informed consent for all study procedures, as did adolescents at T6.

### Participants

Mothers from the community were recruited via local obstetric units and newspapers. Inclusion criteria for the initial study phase were: full-term deliveries, infant weight > 2,500 g, APGAR scores > 7, and good health of the baby during the first three postnatal doctoral exams. Exclusion criteria for mothers were difficulties in German language comprehension, any acute psychiatric disorder, excessive smoking or alcohol consumption, and the use of drugs or medication possibly risking fetal health. At the start of the study, n = 101 mother - infant dyads were enrolled. By the T6 assessment, n = 76 dyads remained, resulting in a retention rate of 75%. Attrition was primarily due to lack of time or interest (9.9%), inability to locate the family (5.9%), maternal death (0.9%), and unspecified reasons (7.9%).

### Birth-related variables

Birth-related factors considered to influence ANS activity included infant sex, birth weight, the pH of the umbilical cord, maternal emotional stress during pregnancy, and any medical complications occurring pre-, peri-, and postnatally. Maternal emotional stress during pregnancy was evaluated with the *Prenatal Emotional Stress Index* (PESI) ^25^, capturing maternal emotional stress across each trimenon. Data on pregnancy and delivery complications were obtained via the *Steinhausen Pre-, Peri-, and Postnatal Score* ^26^, a standardized tool assessing medical complications before, during, and after birth.

### Maternal Bonding Impairment

At T1 (2 weeks postpartum), mothers completed the German version of the *Postpartum Bonding Questionnaire* (PBQ) ^27^, a validated screening instrument designed to identify maternal bonding impairments, assessing dimensions such as impaired bonding, feelings of anger and rejection, anxiety about care, and risk of abuse. Higher scores indicate greater difficulties in maternal-infant bonding. The validity of the PBQ has been confirmed^28^. Following psychometric recommendations, ^27^ we utilized the total PBQ score, which demonstrated satisfactory internal consistency (α = .79).

### Child mental health

Child mental health and behavioral functioning at T5 and T6 was assessed by using the *Strengths and Difficulties Questionnaire* (SDQ) ^29^. The SDQ comprises 25 items covering five subscales: Emotional Symptoms, Conduct Problems, Hyperactivity/Inattention, Peer Problems, and Prosocial Behavior, each consisting of five items. For analytic purposes, the first four subscales are commonly aggregated into two higher-order domains: internalizing problems (Emotional Symptoms and Peer Problems) and externalizing problems (Conduct Problems and Hyperactivity/Inattention), which capture core dimensions of child and adolescent psychopathology. At T5, the SDQ was completed independently by both the child’s mother and preschool teacher. At T6, it was completed by the mother and the child.

### Physiological data recording and preprocessing

ECG data were collected from T1 to T4, and T6. A Colbourn psycho-physiological monitor was used to collect the participants’ ECG during a quiet baseline episode at a frequency of 1000 Hz at T1 to T4. At T6, ECG data were recorded at 1024 Hz using an ECGMove 3 sensor. At T6, participants were seated quietly and instructed to remain at rest during a fixed 5-minute baseline period. At T1–T4, baseline episodes varied in duration, with an average length of 106.11 seconds (SD = 25.84 seconds). To standardize epoch lengths, a 60-second epoch was extracted from each baseline episode from T1 to T4 for further processing, positioned equidistantly from both the beginning and end of the respective episode. For each of the 60-seconds epochs (T1 to T4), the mean HR in beats per minute (bpm) and the root mean square of successive differences (RMSSD) in milliseconds, as measure of vmHRV, were calculated as indices of cardiac autonomic activity using the python toolbox systole ^30^. ECG signals were processed using Systole’s default pipeline, which includes automated detection of R-peaks via a modified Pan-Tompkins algorithm as well as an automated artifact correction procedure based on the subspaces method. The T6 recordings were processed in Kubios HRV 3.0 Premium ^31,32^, to derive RMSSD and HR. In Kubios, peak detection was manually corrected and remaining artifacts were removed. Smoothing priors were selected as detrending method (λ 500) for IBI data.

### Statistical analysis

To characterize children’s age-related trajectories of HR and vmHRV over early development (T1 – T4), we conducted separate linear mixed-effects models for each outcome variable. Children’s age in years at the time of the respective ECG recordings was entered as a continuous predictor. To account for potential sex-specific effects, sex and its interaction with age were additionally included as fixed effects in all models. In both models, a random intercept with a random slope for age was included, accounting for individual variability in age-related trends. The individual slopes were extracted for later analyses. For the timespan from T1 – T4 the full data of n = 90 childern was available and used in the linear mixed models.

To examine the impact of birth-related factors on cardiac outcomes, two separate linear models were applied predicting HR and vmHRV at T1. Each model included infant sex, the infants’ age in days at the assessment day, birth weight, umbilical cord pH, maternal emotional stress during pregnancy (PESI scores), Steinhausen scores, and maternal bonding (PBQ scores) as predictors. To evaluate potential sex-specific effects, infant sex was included as interaction term in both models. For T1 analyses, the full data of n = 101 childern-mother dyads was available.

The impact of birth-related factors on age-related changes in cardiac outcomes was assessed using linear models predicting the age-related slopes of HR and vmHRV extracted from our previous models. Each model used the same set of predictors: infant sex, the infants’ age at the assessment day, birth weight, umbilical cord pH, maternal emotional stress during pregnancy, Steinhausen scores and maternal bonding. The models also accounted for the respective baseline values of HR and vmHRV at two weeks after birth. Infant sex was again included as an interaction term. For the prediction of the age-related slopes, the full data of n = 90 childern-mother dyads was available.

Lastly, to examine the influence of age-related changes in HR and vmHRV on children’s behavioral and emotional health at T5 and T6, individual slopes of age on HR and vmHRV were used as predictors for total scores of the SDQ at T5 and T6. In subsequent analyses, separate models were estimated for the internalizing problems and externalizing problems subscales. In all models, child sex was included to account for potential sex-specific effects. To assess the predictive value of these models, two additional comparison models were estimated for SDQ scores at T6: one including HR and vmHRV measured at T6 only, and another including the change in HR and vmHRV from T4 to T6 as difference scores. For the prediction of the children’s behavioral and emotional health at T5 and T6, the full data of n = 85 and n = 76 childern-mother dyads was available. Sex was effect-coded (−1, 1), and all continuous predictors involved in interaction terms were mean-centered prior to analysis across all models. All statistical analyses were conducted in RStudio^33^, with an alpha error smaller 0.05 considered statistically significant.

## Results

At T1, the sample included n = 101 participants. Full descriptive statistics are provided in *Table 1*. Linear mixed-effects models across early development revealed a significant effect of age on vmHRV (β = .011, *p* < .001) and HR (β = -0.044, *p* < .001), suggesting a general increase in vmHRV (*Figure 1a*) and a decrease in HR (*Figure 1b*) from 2 weeks after birth to 14-months of age. Neither infant sex nor the age × sex interaction significantly predicted vmHRV or HR over time.

**Figure 1.**
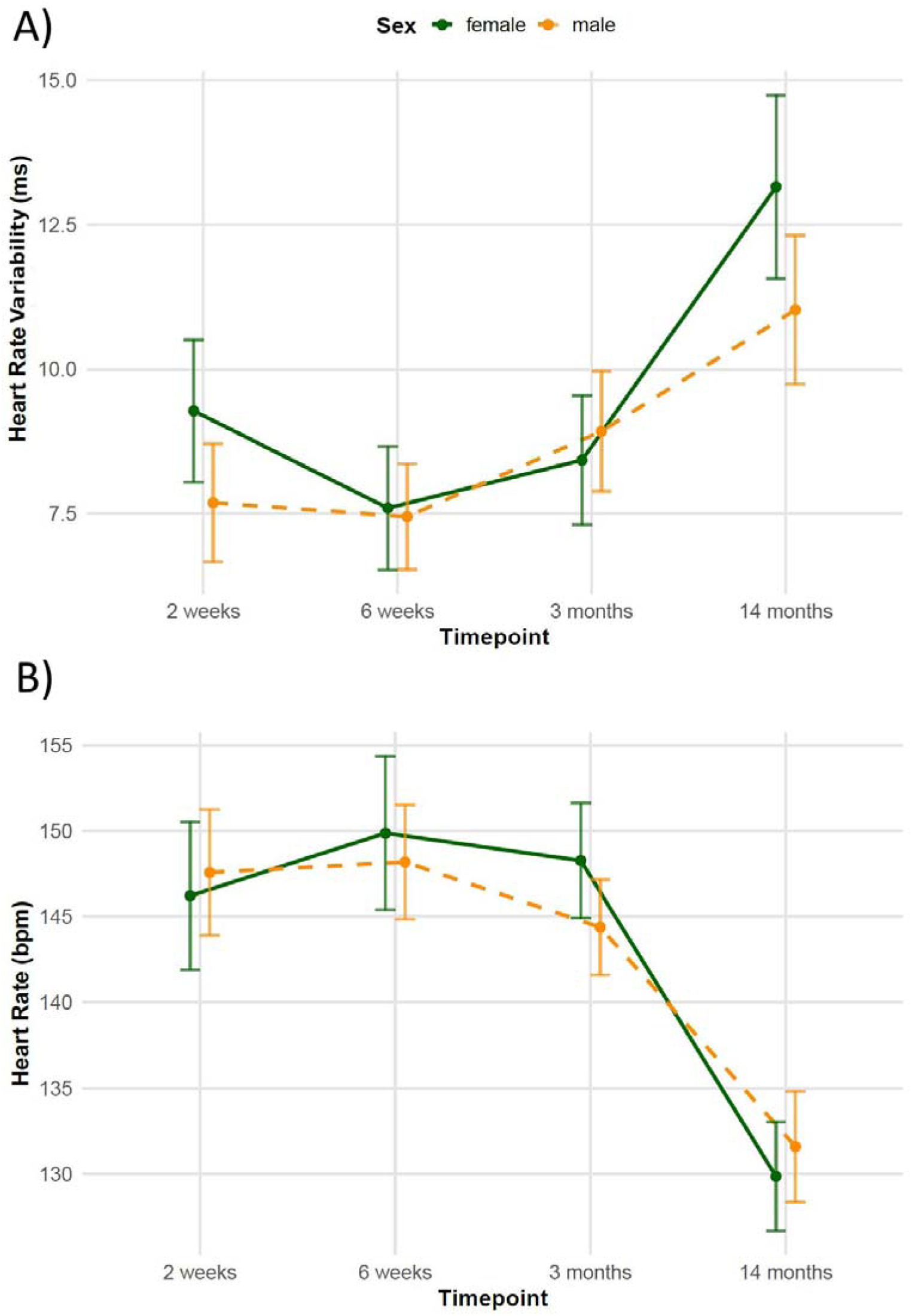
Developmental trajectories of A) vagally-mediated HRV measured by the RMSSD and B) HR between 2 weeks and 14 months of age with vertical bars denoting the 95% CI.

**Table 1.**
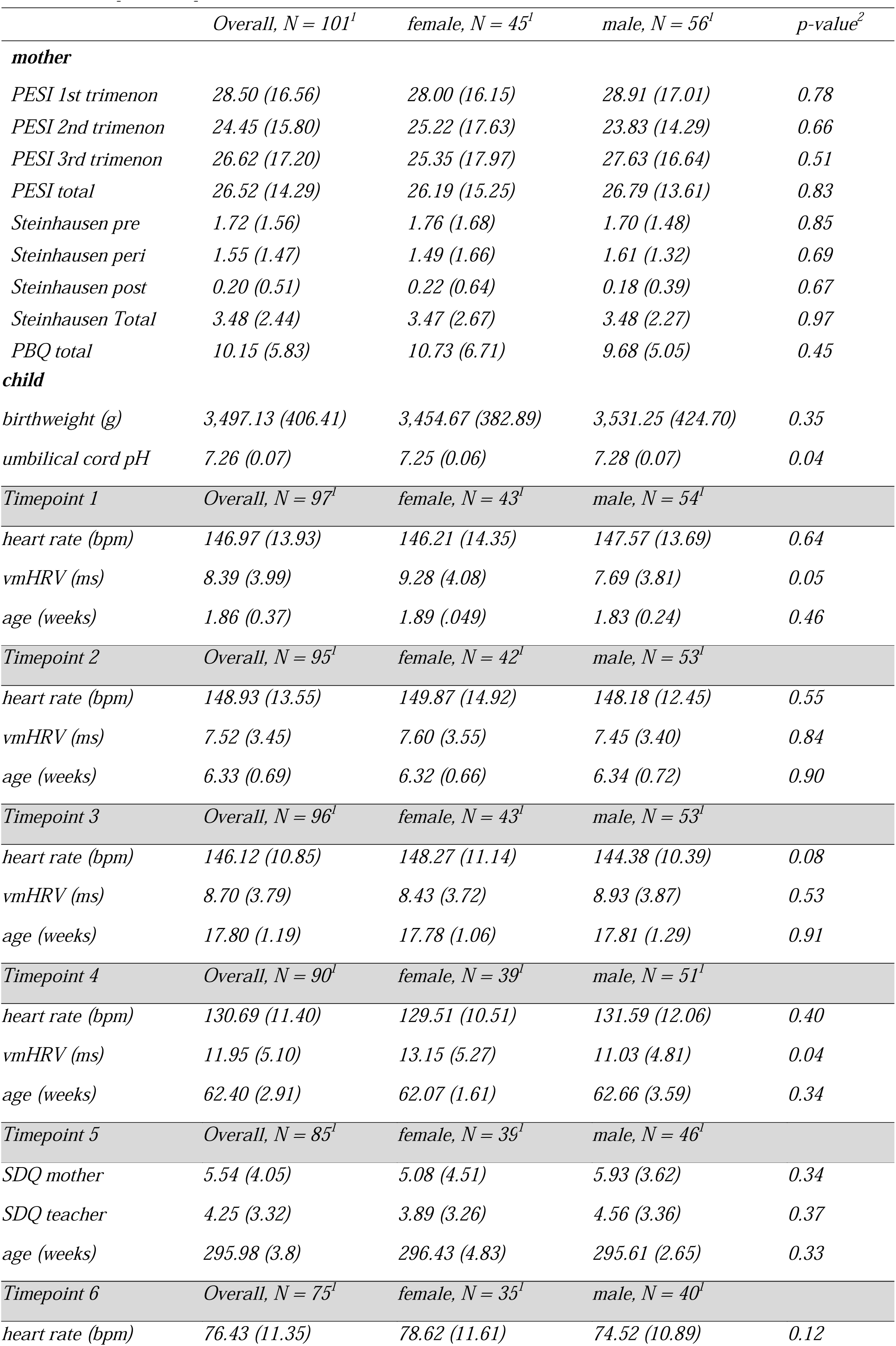
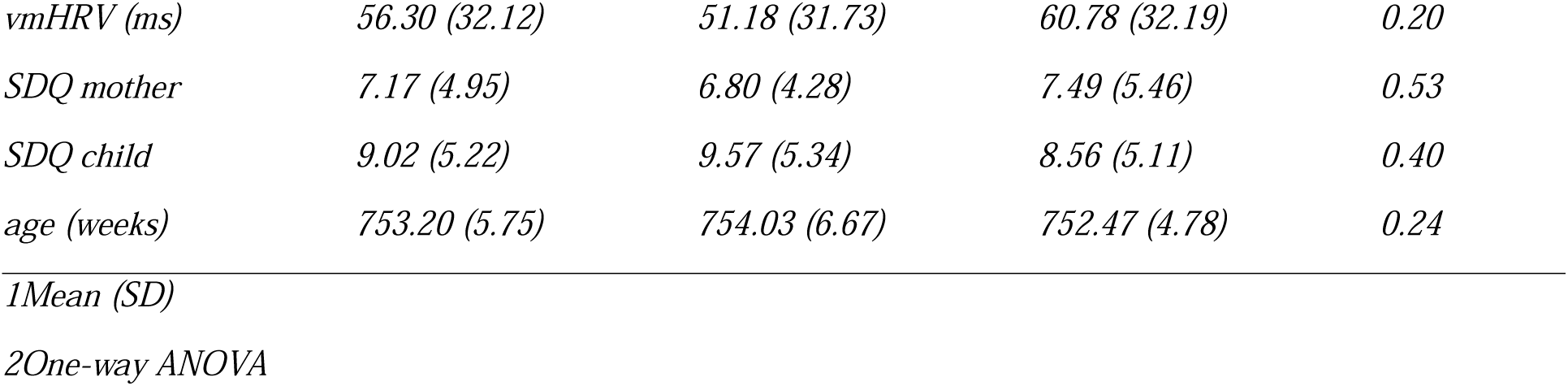
Sample descriptives.

| | Overall, $N = 101^1$ | female, $N = 45^1$ | male, $N = 56^1$ | $p$-value<sup>2</sup> |
| --- | --- | --- | --- | --- |
| **mother** |  |  |  |  |
| *PESI 1st trimenon* | 28.50 (16.56) | 28.00 (16.15) | 28.91 (17.01) | 0.78 |
| *PESI 2nd trimenon* | 24.45 (15.80) | 25.22 (17.63) | 23.83 (14.29) | 0.66 |
| *PESI 3rd trimenon* | 26.62 (17.20) | 25.35 (17.97) | 27.63 (16.64) | 0.51 |
| *PESI total* | 26.52 (14.29) | 26.19 (15.25) | 26.79 (13.61) | 0.83 |
| *Steinhausen pre* | 1.72 (1.56) | 1.76 (1.68) | 1.70 (1.48) | 0.85 |
| *Steinhausen peri* | 1.55 (1.47) | 1.49 (1.66) | 1.61 (1.32) | 0.69 |
| *Steinhausen post* | 0.20 (0.51) | 0.22 (0.64) | 0.18 (0.39) | 0.67 |
| *Steinhausen Total* | 3.48 (2.44) | 3.47 (2.67) | 3.48 (2.27) | 0.97 |
| *PBQ total* | 10.15 (5.83) | 10.73 (6.71) | 9.68 (5.05) | 0.45 |
| **child** |  |  |  |  |
| *birthweight (g)* | 3,497.13 (406.41) | 3,454.67 (382.89) | 3,531.25 (424.70) | 0.35 |
| *umbilical cord pH* | 7.26 (0.07) | 7.25 (0.06) | 7.28 (0.07) | 0.04 |

| *Timepoint 1* | *Overall, N = 97<sup>1</sup>* | *female, N = 43<sup>1</sup>* | *male, N = 54<sup>1</sup>* |  |
| --- | --- | --- | --- | --- |
| *heart rate (bpm)* | 146.97 (13.93) | 146.21 (14.35) | 147.57 (13.69) | 0.64 |
| *vmHRV (ms)* | 8.39 (3.99) | 9.28 (4.08) | 7.69 (3.81) | 0.05 |
| *age (weeks)* | 1.86 (0.37) | 1.89 (.049) | 1.83 (0.24) | 0.46 |
| *Timepoint 2* | *Overall, N = 95<sup>1</sup>* | *female, N = 42<sup>1</sup>* | *male, N = 53<sup>1</sup>* |  |
| *heart rate (bpm)* | 148.93 (13.55) | 149.87 (14.92) | 148.18 (12.45) | 0.55 |
| *vmHRV (ms)* | 7.52 (3.45) | 7.60 (3.55) | 7.45 (3.40) | 0.84 |
| *age (weeks)* | 6.33 (0.69) | 6.32 (0.66) | 6.34 (0.72) | 0.90 |
| *Timepoint 3* | *Overall, N = 96<sup>1</sup>* | *female, N = 43<sup>1</sup>* | *male, N = 53<sup>1</sup>* |  |
| *heart rate (bpm)* | 146.12 (10.85) | 148.27 (11.14) | 144.38 (10.39) | 0.08 |
| *vmHRV (ms)* | 8.70 (3.79) | 8.43 (3.72) | 8.93 (3.87) | 0.53 |
| *age (weeks)* | 17.80 (1.19) | 17.78 (1.06) | 17.81 (1.29) | 0.91 |
| *Timepoint 4* | *Overall, N = 90<sup>1</sup>* | *female, N = 39<sup>1</sup>* | *male, N = 51<sup>1</sup>* |  |
| *heart rate (bpm)* | 130.69 (11.40) | 129.51 (10.51) | 131.59 (12.06) | 0.40 |
| *vmHRV (ms)* | 11.95 (5.10) | 13.15 (5.27) | 11.03 (4.81) | 0.04 |
| *age (weeks)* | 62.40 (2.91) | 62.07 (1.61) | 62.66 (3.59) | 0.34 |
| *Timepoint 5* | *Overall, N = 85<sup>1</sup>* | *female, N = 39<sup>1</sup>* | *male, N = 46<sup>1</sup>* |  |
| *SDQ mother* | 5.54 (4.05) | 5.08 (4.51) | 5.93 (3.62) | 0.34 |
| *SDQ teacher* | 4.25 (3.32) | 3.89 (3.26) | 4.56 (3.36) | 0.37 |
| *age (weeks)* | 295.98 (3.8) | 296.43 (4.83) | 295.61 (2.65) | 0.33 |
| *Timepoint 6* | *Overall, N = 75<sup>1</sup>* | *female, N = 35<sup>1</sup>* | *male, N = 40<sup>1</sup>* |  |
| *heart rate (bpm)* | 76.43 (11.35) | 78.62 (11.61) | 74.52 (10.89) | 0.12 |
| *vmHRV (ms)* | *56.30 (32.12)* | *51.18 (31.73)* | *60.78 (32.19)* | *0.20* |
| *SDQ mother* | *7.17 (4.95)* | *6.80 (4.28)* | *7.49 (5.46)* | *0.53* |
| *SDQ child* | *9.02 (5.22)* | *9.57 (5.34)* | *8.56 (5.11)* | *0.40* |
| *age (weeks)* | *753.20 (5.75)* | *754.03 (6.67)* | *752.47 (4.78)* | *0.24* |

*1Mean (SD)*
*2One-way ANOVA*

### Postnatal Autonomic Function

The linear model predicting vmHRV two weeks after birth explained approximately 25.1% of the variance. Significant predictors included sex, weight, birth complications before and during delivery, umbilical cord pH, as well as the interaction between sex and umbilical cord pH and sex and weight. Specifically, more complications during delivery (β = .80, *p* < .01), and higher umbilical cord pH (β = 19.98, *p* < .01) predicted higher vmHRV at two weeks of age. A significant interaction between sex and umbilical cord pH indicated that this effect was more pronounced in females (*p* < .05). Additionally, higher weight (β = -.004, *p* < .05) and more complications during pregnancy (β = -.60, *p* < .05) predicted lower vmHRV two weeks after birth, with the effect of weight being more pronounced in females (*p* < .05). For detailed results, see *Table 2*. The linear model predicting HR two weeks after birth (see *Table 3*) explained 12.5% of the variance with only complications during delivery emerging as a significant predictor (β = -3.00, *p* < .05).

**Table 2.** Results of linear regression model predicting the infants’ vmHRV two weeks after birth 95 % Confidence Intervall.

| Predictors | Estimates | std.<br>Error | 95 % Confidence<br>Intervall |  | t | p |
| --- | --- | --- | --- | --- | --- | --- |
|  |  |  | Lower | Upper |  |  |
| (Intercept) | 8.57 | 0.40 | 7.77 | 9.38 | 21.22 | <0.001 |
| sex | 1.27 | 0.40 | 0.47 | 2.08 | 3.16 | **0.002** |
| age | -0.29 | 0.20 | -0.68 | 0.10 | -1.50 | 0.139 |
| weight | -0.00 | 0.00 | -0.00 | 0.00 | -1.34 | 0.186 |
| Steinhausen pre | -0.60 | 0.29 | -1.17 | -0.02 | -2.08 | **0.042** |
| Steinhausen post | -0.44 | 0.94 | -2.30 | 1.43 | -0.47 | 0.642 |
| Steinhausen peri | 0.80 | 0.30 | 0.20 | 1.41 | 2.66 | **0.010** |
| umbilical cord pH | 19.98 | 6.89 | 6.23 | 33.73 | 2.90 | **0.005** |
| PESI 1st trimenon | -0.03 | 0.04 | -0.12 | 0.05 | -0.77 | 0.443 |
| PESI 2nd trimenon | 0.01 | 0.05 | -0.09 | 0.12 | 0.27 | 0.790 |
| PESI 3rd trimenon | 0.05 | 0.04 | -0.03 | 0.13 | 1.27 | 0.210 |
| Maternal bonding | 0.06 | 0.07 | -0.09 | 0.21 | 0.83 | 0.408 |
| sex x age | -0.20 | 0.20 | -0.59 | 0.19 | -1.00 | 0.321 |
| sex x weight | -0.00 | 0.00 | -0.00 | -0.00 | -2.14 | **0.037** |
| sex x Steinhausen pre | -0.20 | 0.29 | -0.77 | 0.38 | -0.68 | 0.497 |
| sex x Steinhausen post | 0.52 | 0.94 | -1.35 | 2.38 | 0.55 | 0.583 |
| sex x Steinhausen peri | 0.20 | 0.30 | -0.40 | 0.81 | 0.68 | 0.501 |
| sex x<br>umbilical cord | 21.17 | 6.89 | 7.42 | 34.92 | 3.07 | **0.003** |
| sex x<br>PESI 1st trimenon | 0.00 | 0.04 | -0.08 | 0.09 | 0.08 | 0.936 |
| sex x<br>PESI 2nd trimenon | 0.07 | 0.05 | -0.03 | 0.18 | 1.46 | 0.149 |
| sex x<br>PESI 3rd trimenon | 0.00 | 0.04 | -0.08 | 0.08 | 0.09 | 0.927 |
| sex x<br>Maternal bonding | 0.01 | 0.07 | -0.13 | 0.16 | 0.18 | 0.855 |
| $R^2 / R^2_{\text{adjusted}}$ | | | | | | 0.434 / 0.251 |

**Table 3.** Results of linear regression model predicting the infants’ HR two weeks after birth.

| Predictors | Estimates | std.<br>Error | 95 % Confidence<br>Intervall |  | t | p |
| --- | --- | --- | --- | --- | --- | --- |
|  |  |  | Lower | Upper |  |  |
| (Intercept) | 147.03 | 1.55 | 143.93 | 150.13 | 94.71 | <0.001 |
| sex | -1.03 | 1.55 | -4.13 | 2.07 | -0.66 | 0.509 |
| age | -0.14 | 0.75 | -1.64 | 1.36 | -0.18 | 0.855 |
| weight | 0.01 | 0.00 | -0.00 | 0.02 | 1.87 | 0.066 |
| Steinhausen pre | 1.36 | 1.11 | -0.85 | 3.57 | 1.23 | 0.224 |
| Steinhausen post | 0.52 | 3.59 | -6.66 | 7.70 | 0.15 | 0.885 |
| Steinhausen peri | -3.00 | 1.16 | -5.31 | -0.68 | -2.58 | **0.012** |
| umbilical cord pH | -31.15 | 26.46 | -84.00 | 21.70 | -1.18 | 0.243 |
| PESI 1st trimenon | -0.05 | 0.16 | -0.37 | 0.27 | -0.32 | 0.747 |
| PESI 2nd trimenon | -0.06 | 0.20 | -0.45 | 0.34 | -0.29 | 0.771 |
| PESI 3rd trimenon | 0.12 | 0.15 | -0.18 | 0.43 | 0.81 | 0.423 |
| Maternal bonding | 0.09 | 0.29 | -0.48 | 0.66 | 0.33 | 0.745 |
| sex × age | 0.85 | 0.75 | -0.65 | 2.35 | 1.13 | 0.262 |
| sex × weight | 0.01 | 0.00 | -0.00 | 0.02 | 1.82 | 0.074 |
| sex ×<br>Steinhausen pre | 1.94 | 1.11 | -0.27 | 4.15 | 1.75 | 0.085 |
| sex ×<br>Steinhausen post | -3.27 | 3.59 | -10.45 | 3.90 | -0.91 | 0.366 |
| sex ×<br>Steinhausen peri | 1.71 | 1.16 | -0.61 | 4.02 | 1.47 | 0.146 |
| sex ×<br>umbilical cord pH | -51.82 | 26.46 | -104.67 | 1.03 | -1.96 | 0.054 |
| sex ×<br>PESI 1st trimenon | -0.10 | 0.16 | -0.42 | 0.22 | -0.62 | 0.541 |
| sex ×<br>PESI 2nd trimenon | -0.25 | 0.20 | -0.64 | 0.15 | -1.25 | 0.216 |
| sex ×<br>PESI 3rd trimenon | -0.06 | 0.15 | -0.37 | 0.24 | -0.41 | 0.680 |
| sex ×<br>Maternal bonding | -0.07 | 0.29 | -0.64 | 0.50 | -0.25 | 0.807 |
| $R^2 / R^2$ adjusted | | | | | 0.339 / 0.125 | |

### Autonomic Function during the first 14-months

The linear model predicting the age-related slope of vmHRV during infancy explained 13.9% of the variance with only vmHRV two weeks after birth significantly explaining variance in the vmHRV trajectory, β = .18, *p* < .01. The linear model predicting the age-related slope of HR explained 47.1% of the variance. Analyses revealed that higher HR two weeks after birth significantly predicted a more pronounced decrease in HR over time, β = -.05, *p* < .05. We further identified weight (β = .001, *p* < .05) and complications during delivery (β = -.13, *p* < .05) to significantly predict HR maturation.

### Autonomic and Psychosocial Function at T5 (5 years) and T6 (14 years)

The linear models predicting mean total SDQ scores at T5 showed poor model fit, with adjusted R² values below 0.02 (see *Tables 4 & 5*). Age-related changes in HR and vmHRV from the postnatal period to 14 months did not significantly predict total SDQ scores at T5 (all p > .05), regardless of whether ratings were provided by mothers or preschool teachers. Results from the subscale analyses are presented in Supplementary Tables 1–4.

The linear models predicting total SDQ scores at T6 showed differing model fit depending on the informant. The model based on maternal reports showed better fit (adjusted R² = 0.066) compared to the model based on adolescent self-reports (adjusted R² = –0.013), indicating that the predictors explained more variance in mother-reported than in self-reported SDQ scores (see *Table 6 & 7*). In the maternal-report model, the interaction between sex and HR age slope (β = -2.06, *p* < .01), and the interaction between sex and vmHRV age slope (β = 1.05, *p* < .05) showed statistical significance. Post hoc analyses revealed that flatter age-related decreases in HR were significantly associated with higher SDQ scores in males (β = 3.04, *p* < .05), with a non-significant association for females. Similarly, a steeper increase in vmHRV was significantly associated with lower SDQ scores only in males, β = – 1.53, *p* < .05. No predictors were significantly associated with self-reported SDQ scores at age 14 (all p > .05). For the results of the subscale analyses, see Supplementary Tables 5-8. For the results of the comparison models, see Supplementary Tables 9-12.

**Table 4.** Results of linear regression model predicting maternal-reported total SDQ scores at the age of 5 years.

| Predictors | Estimates | std.<br>Error | 95 % Confidence<br>Intervall |  | t | p |
| --- | --- | --- | --- | --- | --- | --- |
|  |  |  | Lower | Upper |  |  |
| (Intercept) | 5.42 | 0.44 | 4.54 | 6.30 | 12.30 | <0.001 |
| sex | -0.37 | 0.44 | -1.25 | 0.50 | -0.85 | 0.398 |
| HR age slope | 0.17 | 0.55 | -0.93 | 1.26 | 0.31 | 0.759 |
| vmHRV age slope | -0.49 | 0.34 | -1.17 | 0.18 | -1.45 | 0.151 |
| sex * HR age slope | -0.92 | 0.54 | -2.01 | 0.17 | -1.68 | 0.097 |
| sex * vmHRV age slope | 0.32 | 0.34 | -0.35 | 1.00 | 0.96 | 0.340 |
| $R^2$ / $R^2$ adjusted | | | | | | 0.072 / 0.013 |

**Table 5.** Results of linear regression model predicting preschool teacher-reported total SDQ scores at the age of 5 years.

| Predictors | Estimates | std.<br>Error | 95 % Confidence<br>Intervall |  | t | p |
| --- | --- | --- | --- | --- | --- | --- |
|  |  |  | Lower | Upper |  |  |
| (Intercept) | 4.23 | 0.3831 | 3.46 | 4.99 | 11.05 | <0.001 |
| sex | -0.37 | 0.38 | -1.14 | 0.39 | -0.97 | 0.334 |
| HR age slope | -0.30 | 0.46 | -1.23 | 0.62 | -0.66 | 0.515 |
| vmHRV age slope | 0.24 | 0.29 | -0.35 | 0.82 | 0.81 | 0.419 |
| sex * HR age slope | -0.14 | 0.46 | -1.06 | 0.77 | -0.31 | 0.757 |
| sex * vmHRV age slope | -0.18 | 0.29 | -0.76 | 0.40 | -0.63 | 0.533 |
| $R^2$ / $R^2$ adjusted | | | | | | 0.036 / -0.030 |

**Table 6.** Results of linear regression model predicting maternal-reported total SDQ scores at the age of 14 years.

| *Predictors* | *Estimates* | *std.*<br>*Error* | 95 % Confidence<br>Intervall |  | *t* | *p* |
| --- | --- | --- | --- | --- | --- | --- |
|  |  |  | *Lower* | *Upper* |  |  |
| *(Intercept)* | 6.88 | 0.56 | 5.76 | 7.99 | 12.30 | <0.001 |
| *sex* | -0.19 | 0.56 | -1.30 | 0.93 | -0.33 | 0.739 |
| *HR age slope* | 0.98 | 0.73 | -0.49 | 2.44 | 1.33 | 0.188 |
| *vmHRV age slope* | -0.48 | 0.42 | -1.31 | 0.36 | -1.14 | 0.257 |
| *sex \* HR age slope* | -2.06 | 0.73 | -3.53 | -0.60 | -2.81 | **0.006** |
| *sex \* vmHRV age slope* | 1.05 | 0.42 | 0.21 | 1.88 | 2.50 | **0.015** |
| *R<sup>2</sup> / R<sup>2</sup> adjusted* | 0.128 / 0.066 |  |  |  |  |  |

**Table 7.** Results of linear regression model predicting self-reported total SDQ scores at the age of 14 years.

| *Predictors* | *Estimates* | *std.*<br>*Error* | 95 % Confidence<br>Intervall |  | *t* | *p* |
| --- | --- | --- | --- | --- | --- | --- |
|  |  |  | *Lower* | *Upper* |  |  |
| *(Intercept)* | 9.03 | 0.61 | 7.81 | 10.26 | 14.69 | <0.001 |
| *sex* | 0.58 | 0.61 | -0.64 | 1.81 | 0.95 | 0.347 |
| *HR age slope* | 0.76 | 0.81 | -0.85 | 2.37 | 0.94 | 0.352 |
| *vmHRV age slope* | -0.34 | 0.46 | -1.26 | 0.57 | -0.75 | 0.458 |
| *sex \* HR age slope* | -0.45 | 0.81 | -2.06 | 1.16 | -0.56 | 0.577 |
| *sex \* vmHRV age slope* | 0.71 | 0.46 | -0.21 | 1.63 | 1.55 | 0.127 |
| *R<sup>2</sup> / R<sup>2</sup> adjusted* | 0.052 / -0.013 |  |  |  |  |  |

## Discussion

Developmental neurovisceral models propose that ANS maturation is tightly interconnected with brain development and the emergence of adaptive psychological functions, ^14^ with adverse early life experiences ^12^ impacting this brain-body synchrony and potentially leading to maladaptive cognitive and emotional trajectories. In a first and extensive longitudinal study, we aimed to investigate the association of early ANS maturation with psychosocial outcomes.

Highlighting key aspects of early autonomic maturation, we found significant age-related changes in both vmHRV and HR. Consistent with earlier findings ^34^, we observed an initial descriptive increase in HR from two to six weeks of age, followed by a gradual decline. VmHRV inversely mirrored this pattern with a distinct negative peak at six weeks followed by a constant increase. This pattern aligns with prior research ^35–38^, reporting an early postnatal decline in vmHRV. While these findings suggest a period of heightened autonomic reorganization during the first weeks of life, reflecting dynamic adaptations in vagal regulation, further research is warranted, investigating mechanisms underlying these early autonomic adaptations, in the interplay of HR and vmHRV, that seem independent of sex.

Our findings further suggest that vmHRV two weeks after birth is affected by multiple birth-related factors. Specifically, higher umbilical pH was linked to greater vmHRV, with stronger effects in females. Similarly, we found that only in females, higher birth weight was associated with lower vmHRV. Previous studies investigating the relationship between birth weight and vmHRV shortly after birth, reported mixed findings. While some studies observed no significant association ^39,40^, others reported positive correlations. ^37,41^ In respective studies, these effects, however, were mainly driven by subgroups of infants with very low birth weight, exhibiting significantly reduced vmHRV compared to newborns with normal birth weight. While in the present study, the first vmHRV recording was conducted two weeks after birth, the studies mentioned above measured vmHRV within a few hours to several days postnatally. As systematic changes in vmHRV have been shown to occur as early as six hours after birth, ^42^ contrasting results might arise from differences in the timing of recordings. As such, birth weight may especially in female newborns influence vagal adaptation in a way that its effects only become apparent at later stages of neonatal development. Furthermore, consistent with our findings, previous research in humans ^43–46^ and animal models ^47^ has shown that fetal acidosis, reflected by reduced umbilical cord pH due to inadequate oxygenation, is associated with alterations in fetal vmHRV. Accordingly, the observed positive correlation between umbilical cord pH and higher vmHRV two weeks postnatally suggests that oxygenation levels during birth may critically influence early maturation of the ANS.

Complications during pregnancy and delivery emerged as further predictors of vmHRV two weeks postpartum. Specifically, a greater number of pregnancy complications predicted lower vmHRV, while more problematic events during delivery were associated with higher vmHRV. These findings align with previous research demonstrating the adverse effects of multiple factors during pregnancy such as maternal substance use ^48^ or mental health challenges ^48,49^ on the newborns’ ANS. However, our observation of a positive association between vmHRV and delivery complications contrasts with earlier studies, which found no association between delivery complications and vmHRV immediately after birth ^42,50^. It has to be noted that the Steinhausen Perinatal Score used here also includes the occurrence of non-spontaneous delivery modes. Moehler et al., ^13^ analyzing the same dataset, found that infants delivered via vacuum extraction exhibited higher vmHRV and lower HR compared to those born spontaneously or via cesarean section (with no significant differences between spontaneous and cesarean deliveries). As such, our results suggest that the positive association between vmHRV and the Steinhausen Perinatal Score may be at least partially driven by the influence of non-spontaneous delivery modes. These findings collectively suggest that delivery mode as well as further events during delivery may exert differential effects on ANS development depending on postnatal age.

While results showed a significant effect of birth-related factors on vmHRV at two weeks postpartum, no such effect was found on the longitudinal changes in vmHRV during infancy. This finding suggests that during the first year of life, the ANS undergoes substantial developmental changes, independent of early birth-related factors. Supporting this notion, it has been demonstrated that although preterm infants exhibit deficits in ANS activity immediately after birth compared to full-term infants, by two years of age, they achieve comparable maturation of the ANS. ^51^ However, in contrast to these results, we found higher vmHRV at two weeks postpartum to predict steeper increases in vmHRV across the first 14 months of life. This suggests that early vagal tone may not merely reflect transient perinatal status but instead index the functional quality of initial autonomic organization, which in turn may scaffold subsequent parasympathetic development. Similarly, HR at two weeks significantly predicted the developmental trajectory of HR over time with higher HR being associated with a steeper decline in HR. Furthermore, we found higher birth weight as well as more delivery complications to predict HR trajectories. Specifically, higher birth weight was linked to flatter decreases in HR while an opposite association was found for delivery complications.

We further investigated whether early developmental ANS trajectories between the age of two weeks and 14 months predict emotional and behavioral outcomes at the age of 5 and 14 years. The present findings offer novel longitudinal evidence that early developmental trajectories of autonomic function, specifically, age-related changes in HR and vmHRV from 2 weeks to 14 months, are associated with psychosocial functioning more than a decade later. While these autonomic indices did not predict maternal- or teacher-rated SDQ scores at age 5, flatter decreases in HR and less steep increases in vmHRV significantly predicted higher SDQ scores at age 14, as reported by mothers. Notably, these early developmental slopes predicted SDQ outcomes more strongly than contemporaneous HR and HRV values measured at age 14 or the change in both measures from T4 to T6 (see Supplementary Tables 9 - 12), underscoring the value of longitudinal indicators over static cross-sectional assessments. This effect was specific to males and predominantly driven by externalizing problems (see Supplementary Table 5). These findings align with a growing body of research demonstrating that subtle neurobiological markers can prospectively forecast the emergence of psychiatric symptoms during adolescence, even in the absence of overt behavioral problems earlier in life. ^52–54^ The absence of associations at age 5 may reflect a developmental lag, wherein early autonomic deviations remain behaviorally latent until the increasing demands of adolescence interact with neurobiological vulnerabilities to unmask underlying regulatory deficits. This pattern has also been observed in large-scale longitudinal imaging studies, which identify similar delayed emergence of behavioral risk associated with early neurobiological variation ^55–57^. The observed sex-specific effects, with early autonomic dysmaturation predicting psychosocial problems predominantly in males, are consistent with broader literature on sex differences in neurodevelopmental trajectories and susceptibility to behavioral dysregulation ^58^. These findings emphasize the importance of modeling early-life physiological development across time, rather than relying solely on concurrent measurements, to improve prediction of adolescent mental health outcomes.

Despite its strengths, particularly the long within-subject timespan, several limitations should be considered. First, the study focused primarily on pregnancy- and delivery-related factors, where as postnatal influences known to affect infant autonomic functioning such as maternal stress, ^59^ emotional health, ^60^ and early mother-infant interactions ^61–63^ were not comprehensively captured. These factors may have contributed to variability in ANS trajectories beyond the scope of the present measures. Second, ECG recordings across the study were relatively sparse, resulting in substantial gaps between recordings, particularly during critical developmental periods between 3 to 14 months and 5 to 14 years of age. This limits the temporal resolution of ANS developmental modeling. More frequent assessments during these periods would allow for finer-grained characterization of autonomic maturation and its links to later psychosocial outcomes. Additionally, the absence of systematic assessment of life events between ages 5 and 14 presents another limitation. This period is typically characterized by substantial environmental and psychosocial changes known to influence autonomic regulation and mental health,. ^64–66^ which were not captured here. Lastly, only short-term resting-state ECG recordings were used in this study. Long-term assessments could offer a more comprehensive understanding of the diurnal dynamicas of vagal activity which have been linked to mental health conditions characterized by emotional and social impairments. ^67,68^ Future longitudinal work integrating extended physiological monitoring would therefore improve understanding of how early experiences shape long-term vagal regulation and its relation to mental health.

Taken together, this study underscores that early ANS development is meaningfully associated with later psychosocial functioning. While birth-related and maternal factors significantly influenced cardiac activity at two weeks of age, they did not shape subsequent parasympathetic developmental trajectories, suggesting a high degree of postnatal plasticity in autonomic maturation. Individual differences in early ANS trajectories themselves showed prospective associations with psychosocial outcomes even years after the last ECG recording, reinforcing the role of ANS maturation as a critical antecedent for emotion regulation and social behavior. These effects were more pronounced in males, highlighting potential sex-specific pathways in neurodevelopmental risk. Conceptually, these findings support models linking early autonomic regulation to the organization of broader emotion-regulatory and social-cognitive systems, and emphasize the importance of capturing developmental change rather than static levels of physiological function. More broadly, they suggest that early-life autonomic trajectories may help identify sensitive periods for intervention, during which targeted support could optimize long-term emotional and behavioral adaptation.

## Data Availability

The datasets generated and analyzed, along with the code used in this study, are available from the corresponding author upon reasonable request.

## Contributions

Conceptualization, L.F., A.F., E.M., M.K., and J.K. Methodology, M.S., L.F., A.F., M.K., and J.K.; Investigation, A.F., L.F., and E.M.; Data Curation, L.F. and A.F.; Formal Analysis, M.S.; Writing – Original Draft, M.S.; Writing – Review & Editing, all authors; Funding Acquisition, M.K. and J.K.; Resources, M.K. and J.K.; Supervision, M.K. and J.K.

## Acknowledgments

Julian Koenig acknowledges financial support for the Mapping Autonomic Neural Interaction and Control (MANIAC) Emerging Group by the University of Cologne Excellent Research Support Program.

## Disclosure

All authors declare that they have no biomedical financial interests or other conflicts of interest, financial or otherwise, to disclose.

## Notes

### Competing Interest Statement

The authors have declared no competing interest.

