## Supplementary Material for "Postnatal Autonomic Development Predicts Adolescent Psychosocial Outcome"

| *Supplementary Table 1.* Results of linear regression model predicting maternal-reported SDQ externalizing subscale scores at the age of 5 years | | | | | | | |
| --- | --- | --- | --- | --- | --- | --- | --- |
|  |  |  |  | *95 % Confidence Intervall* | |  |  |
| *Predictors* |  | *Estimates* | *std. Error* | *Lower* | *Upper* | *t* | *p* |
|  | *(Intercept)* | *4.30* | *0.42* | *3.46* | *5.14* | *10.24* | *<0.001* |
| *sex* |  | *-0.48* | *0.42* | *-1.32* | *0.35* | *-1.15* | *0.253* |
| *HR age slope* |  | *0.02* | *0.52* | *-1.02* | *1.06* | *0.04* | *0.970* |
| *vmHRV age slope* |  | *-0.08* | *0.32* | *-0.73* | *0.56* | *-0.25* | *0.803* |
| *sex * HR age slope* |  | *-1.09* | *0.52* | *-2.14* | *-0.05* | *-2.09* | ***0.040*** |
| *sex * vmHRV age slope* |  | *0.61* | *0.32* | *-0.04* | *1.25* | *1.87* | *0.065* |
| *R^2^ / R^2^ adjusted* |  | | *0.077 / 0.018* | | | | |

| *Supplementary Table 2.* Results of linear regression model predicting maternal-reported SDQ internalizing subscale scores at the age of 5 years | | | | | | | |
| --- | --- | --- | --- | --- | --- | --- | --- |
|  |  |  |  | *95 % Confidence Intervall* | |  |  |
| *Predictors* |  | *Estimates* | *std. Error* | *Lower* | *Upper* | *t* | *p* |
|  | *(Intercept)* | *1.72* | *0.23* | *1.27* | *2.17* | *7.65* | *<0.001* |
| *sex* |  | *0.24* | *0.23* | *-0.21* | *0.69* | *1.06* | *0.295* |
| *HR age slope* |  | *0.02* | *0.28* | *-0.54* | *0.58* | *0.08* | *0.940* |
| *vmHRV age slope* |  | *-0.19* | *0.17* | *-0.53* | *0.16* | *-1.07* | *0.289* |
| *sex * HR age slope* |  | *-0.09* | *0.28* | *-0.65* | *0.47* | *-0.31* | *0.755* |
| *sex * vmHRV age slope* |  | *-0.09* | *0.17* | *-0.44* | *0.25* | *-0.53* | *0.600* |
| *R^2^ / R^2^ adjusted* |  | | *0.039 / -0.020* | | | | |

| *Supplementary Table 3.* Results of linear regression model predicting preschool teacher-reported SDQ externalizing subscale scores at the age of 5 years | | | | | | | |
| --- | --- | --- | --- | --- | --- | --- | --- |
|  |  |  |  | *95 % Confidence Intervall* | |  |  |
| *Predictors* |  | *Estimates* | *std. Error* | *Lower* | *Upper* | *t* | *p* |
|  | *(Intercept)* | *3.65* | *0.48* | *2.70* | *4.60* | *7.68* | *<0.001* |
| *sex* |  | *-0.81* | *0.48* | *-1.75* | *0.14* | *-1.69* | *0.095* |
| *HR age slope* |  | *-0.12* | *0.57* | *-1.27* | *1.02* | *-0.22* | *0.830* |
| *vmHRV age slope* |  | *-0.23* | *0.36* | *-0.96* | *0.49* | *-0.64* | *0.525* |
| *sex * HR age slope* |  | *0.23* | *0.57* | *-0.92* | *1.37* | *0.39* | *0.695* |
| *sex * vmHRV age slope* |  | *-0.38* | *0.36* | *-1.10* | *0.35* | *-1.03* | *0.307* |
| *R^2^ / R^2^ adjusted* |  | | *0.063 / -0.001* | | | | |

| *Supplementary Table 4.* Results of linear regression model predicting preschool teacher-reported SDQ internalizing subscale scores at the age of 5 years | | | | | | | |
| --- | --- | --- | --- | --- | --- | --- | --- |
|  |  |  |  | *95 % Confidence Intervall* | |  |  |
| *Predictors* |  | *Estimates* | *std. Error* | *Lower* | *Upper* | *t* | *p* |
|  | *(Intercept)* | *1.51* | *0.22* | *1.07* | *1.95* | *6.82* | *<0.001* |
| *sex* |  | *0.19* | *0.22* | *-0.24* | *0.64* | *0.90* | *0.369* |
| *HR age slope* |  | *-0.13* | *0.27* | *-0.66* | *0.41* | *-0.47* | *0.639* |
| *vmHRV age slope* |  | *0.19* | *0.17* | *-0.15* | *0.53* | *1.12* | *0.267* |
| *sex * HR age slope* |  | *-0.19* | *0.27* | *-0.72* | *0.34* | *-0.71* | *0.479* |
| *sex * vmHRV age slope* |  | *0.05* | *0.17* | *-0.29* | *0.39* | *0.29* | *0.772* |
| *R^2^ / R^2^ adjusted* |  | | *0.039 / -0.026* | | | | |

| *Supplementary Table 5.* Results of linear regression model predicting maternal-reported SDQ externalizing subscale scores at the age of 14 years | | | | | | | |
| --- | --- | --- | --- | --- | --- | --- | --- |
|  |  |  |  | *95 % Confidence Intervall* | |  |  |
| *Predictors* |  | *Estimates* | *std. Error* | *Lower* | *Upper* | *t* | *p* |
|  | *(Intercept)* | *4.68* | *0.48* | *3.72* | *5.64* | *9.73* | *<0.001* |
| *sex* |  | *0.03* | *0.48* | *-0.93* | *0.99* | *0.07* | *0.946* |
| *HR age slope* |  | *0.00* | *0.63* | *-1.25* | *1.26* | *0.01* | *0.994* |
| *vmHRV age slope* |  | *0.25* | *0.36* | *-0.47* | *0.97* | *0.69* | *0.494* |
| *sex * HR age slope* |  | *-1.56* | *0.63* | *-2.82* | *-0.30* | *-2.47* | ***0.016*** |
| *sex * vmHRV age slope* |  | *0.93* | *0.36* | *0.21* | *1.65* | *2.59* | ***0.012*** |
| *R^2^ / R^2^ adjusted* |  | | *0.115 / 0.052* | | | | |

| *Supplementary Table 6.* Results of linear regression model predicting maternal-reported SDQ internalizing subscale scores at the age of 14 years | | | | | | | |
| --- | --- | --- | --- | --- | --- | --- | --- |
|  |  |  |  | *95 % Confidence Intervall* | |  |  |
| *Predictors* |  | *Estimates* | *std. Error* | *Lower* | *Upper* | *t* | *p* |
|  | *(Intercept)* | *2.75* | *0.33* | *2.09* | *3.42* | *8.24* | *<0.001* |
| *sex* |  | *-0.23* | *0.33* | *-0.90* | *0.43* | *-0.70* | *0.485* |
| *HR age slope* |  | *0.64* | *0.44* | *-0.23* | *1.52* | *1.47* | *0.147* |
| *vmHRV age slope* |  | *-0.39* | *0.25* | *-0.89* | *0.11* | *-1.56* | *0.124* |
| *sex * HR age slope* |  | *-0.85* | *0.44* | *-1.72* | *0.02* | *-1.94* | *0.057* |
| *sex * vmHRV age slope* |  | *0.36* | *0.25* | *-0.14* | *0.86* | *1.43* | *0.156* |
| *R^2^ / R^2^ adjusted* |  | | *0.089 / 0.024* | | | | |

| *Supplementary Table 8.* Results of linear regression model predicting self-reported SDQ internalizing subscale scores at the age of 14 years | | | | | | | |
| --- | --- | --- | --- | --- | --- | --- | --- |
|  |  |  |  | *95 % Confidence Intervall* | |  |  |
| *Predictors* |  | *Estimates* | *std. Error* | *Lower* | *Upper* | *t* | *p* |
|  | *(Intercept)* | *4.24* | *0.34* | *3.55* | *4.92* | *12.40* | *<0.001* |
| *sex* |  | *0.37* | *0.34* | *-0.31* | *1.06* | *1.10* | *0.277* |
| *HR age slope* |  | *1.05* | *0.45* | *0.15* | *1.94* | *2.34* | *0.022* |
| *vmHRV age slope* |  | *-0.35* | *0.26* | *-0.86* | *0.16* | *-1.38* | *0.172* |
| *sex * HR age slope* |  | *-0.21* | *0.45* | *-1.10* | *0.69* | *-0.46* | *0.647* |
| *sex * vmHRV age slope* |  | *0.36* | *0.26* | *-0.15* | *0.87* | *1.41* | *0.164* |
| *R^2^ / R^2^ adjusted* |  | | *0.100 / 0.036* | | | | |

| *Supplementary Table 7.* Results of linear regression model predicting self-reported SDQ externalizing subscale scores at the age of 14 years | | | | | | | |
| --- | --- | --- | --- | --- | --- | --- | --- |
|  |  |  |  | *95 % Confidence Intervall* | |  |  |
| *Predictors* |  | *Estimates* | *std. Error* | *Lower* | *Upper* | *t* | *p* |
|  | *(Intercept)* | *6.40* | *0.53* | *5.33* | *7.46* | *11.97* | *<0.001* |
| *sex* |  | *0.30* | *0.53* | *-0.76* | *1.37* | *0.57* | *0.571* |
| *HR age slope* |  | *-0.17* | *0.70* | *-1.57* | *1.23* | *-0.24* | *0.809* |
| *vmHRV age slope* |  | *0.01* | *0.40* | *-0.79* | *0.81* | *0.02* | *0.984* |
| *sex * HR age slope* |  | *-0.34* | *0.70* | *-1.74* | *1.06* | *-0.48* | *0.632* |
| *sex * vmHRV age slope* |  | *0.60* | *0.40* | *-0.20* | *1.40* | *1.50* | *0.138* |
| *R^2^ / R^2^ adjusted* |  | | *0.042 / -0.026* | | | | |

| *Supplementary Table 9.* HR and vmHRV at the age of 14 years predicting maternal-reported SDQ scores at the age of 14 years | | | | | | | | |
| --- | --- | --- | --- | --- | --- | --- | --- | --- |
|  |  |  |  | *95 % Confidence Intervall* | | |  |  |
| *Predictors* |  | *Estimates* | *std. Error* | *Lower* | *Upper* | | *t* | *p* |
|  | *(Intercept)* | *7.23* | *0.60* | *6.03* | *8.42* | *12.09* | | *<0.001* |
| *sex* |  | *-0.40* | *0.60* | *-1.59* | *0.79* | *-0.67* | | *0.508* |
| *HR* |  | *0.06* | *0.07* | *-0.07* | *0.19* | *0.91* | | *0.367* |
| *vmHRV* |  | *0.01* | *0.02* | *-0.03* | *0.06* | *0.51* | | *0.610* |
| *sex * HR* |  | *-0.06* | *0.07* | *-0.19* | *0.07* | *-0.89* | | *0.379* |
| *sex * vmHRV* |  | *-0.01* | *0.02* | *-0.05* | *0.04* | *-0.25* | | *0.802* |
| *R^2^ / R^2^ adjusted* |  | | *0.028 / -0.043* | | | | | |

| *Supplementary Table 10.* HR and vmHRV at the age of 14 years predicting self-reported SDQ scores at the age of 14 years | | | | | | | |
| --- | --- | --- | --- | --- | --- | --- | --- |
|  |  |  |  | *95 % Confidence Intervall* | |  |  |
| *Predictors* |  | *Estimates* | *std. Error* | *Lower* | *Upper* | *t* | *p* |
|  | *(Intercept)* | *9.12* | *0.63* | *7.87* | *10.38* | *14.49* | *<0.001* |
| *sex* |  | *0.44* | *0.63* | *-0.81* | *1.70* | *0.70* | *0.485* |
| *HR* |  | *-0.04* | *0.07* | *-0.18* | *0.10* | *-0.53* | *0.600* |
| *vmHRV* |  | *-0.03* | *0.02* | *-0.08* | *0.02* | *-1.15* | *0.254* |
| *sex * HR* |  | *-0.02* | *0.07* | *-0.16* | *0.12* | *-0.26* | *0.796* |
| *sex * vmHRV* |  | *0.00* | *0.02* | *-0.05* | *0.05* | *0.12* | *0.902* |
| *R^2^ / R^2^ adjusted* |  | | *0.033 / -0.038* | | | | |

| *Supplementary Table 11.* HR and vmHRV changes from T4 to T6 predicting maternal-reported SDQ scores at the age of 14 years | | | | | | | |
| --- | --- | --- | --- | --- | --- | --- | --- |
|  |  |  |  | *95 % Confidence Intervall* | |  |  |
| *Predictors* |  | *Estimates* | *std. Error* | *Lower* | *Upper* | *t* | *p* |
|  | *(Intercept)* | *7.29* | *0.65* | *5.99* | *8.58* | *11.24* | *<0.001* |
| *sex* |  | *-0.13* | *0.65* | *-1.43* | *1.17* | *-0.20* | *0.840* |
| *HR* |  | *0.02* | *0.05* | *-0.08* | *0.13* | *0.45* | *0.653* |
| *vmHRV* |  | *0.00* | *0.02* | *-0.04* | *0.04* | *0.06* | *0.951* |
| *sex * HR* |  | *0.02* | *0.05* | *-0.09* | *0.13* | *0.36* | *0.720* |
| *sex * vmHRV* |  | *0.01* | *0.02* | *-0.04* | *0.05* | *0.33* | *0.743* |
| *R^2^ / R^2^ adjusted* |  | | *0.006 / -0.076* | | | | |

| *Supplementary Table 12.* HR and vmHRVchange from T5 to T6 predicting self-reported SDQ scores at the age of 14 years | | | | | | | |
| --- | --- | --- | --- | --- | --- | --- | --- |
|  |  |  |  | *95 % Confidence Intervall* | |  |  |
| *Predictors* |  | *Estimates* | *std. Error* | *Lower* | *Upper* | *t* | *p* |
|  | *(Intercept)* | *9.17* | *0.67* | *7.82* | *10.52* | *13.62* | *<0.001* |
| *sex* |  | *0.57* | *0.67* | *-0.78* | *1.92* | *0.85* | *0.401* |
| *HR* |  | *0.02* | *0.06* | *-0.09* | *0.13* | *0.37* | *0.711* |
| *vmHRV* |  | *-0.02* | *0.02* | *-0.06* | *0.03* | *-0.86* | *0.392* |
| *sex * HR* |  | *0.09* | *0.06* | *-0.02* | *0.20* | *1.62* | *0.112* |
| *sex * vmHRV* |  | *0.02* | *0.02* | *-0.02* | *0.07* | *1.05* | *0.296* |
| *R^2^ / R^2^ adjusted* |  | | *0.074 / -0.002* | | | | |
